# A global *Brassica* pest and a sympatric cryptic ally, *Plutella australiana* (Lepidoptera: Plutellidae), show strong divergence despite the capacity to hybridize

**DOI:** 10.1101/189266

**Authors:** Kym D. Perry, Gregory J. Baker, Kevin J. Powis, Joanne K. Kent, Christopher M. Ward, Simon W. Baxter

**Author notes:** Corresponding author: Kym D. Perry.

## Abstract

The diamondback moth, *Plutella xylostella*, has been intensively studied due to its ability to evolve insecticide resistance and status as the world’s most destructive pest of brassicaceous crops. The surprise discovery of a cryptic ally, *Plutella australiana* Landry & Hebert, with apparent endemism to Australia, immediately raised questions regarding the extent of ecological and genetic diversity between these two species, whether gene flow could occur, and ultimately if specific management was required. Here, we show that despite sympatric distributions and the capacity to hybridize in controlled laboratory experiments, striking differences in genetic and phenotypic traits exist that are consistent with contrasting colonization histories and reproductive isolation after secondary contact. Almost 1500 *Plutella* individuals were collected from wild and cultivated brassicaceous plants at 75 locations throughout Australia. *Plutella australiana* was commonly found on all *Brassica* host types sampled except commercial vegetables, which are routinely sprayed with insecticide. Bioassays using four commonly-used insecticides found that *P. australiana* was 19-306 fold more susceptible than *P. xylostella*. Genome-wide SNPs derived from RADseq revealed substantially higher levels of genetic diversity across *P. australiana* compared with *P. xylostella* nuclear genomes, yet both species showed limited variation in mtDNA. Infection with a single *Wolbachia* subgroup B strain was fixed in *P. australiana*, suggesting that a selective sweep contributed to low mtDNA diversity, while a subgroup A strain infected just 1.5 % of *P. xylostella*. Although *P. australiana* is a potential pest of brassica crops, it is of secondary importance to *P. xylostella*.

## Introduction

Cryptic species share morphological traits, yet can show remarkable diversity in aspects of their ecology, behaviour, and at the level of the genome. They exist across metazoan taxa (Pfenninger & Schwenk, 2007), including globally important arthropod pest taxa, such as white-flies (De Barro, Liu, Boykin, & Dinsdale, 2011), mosquito vectors (Coetzee et al., 2013), fruit flies (Hendrichs, Teresa Vera, De Meyer, & Clarke, 2015), thrips (Jacobson, Nault, Vargo, & Kennedy, 2016; Rugman-Jones, Hoddle, & Stouthamer, 2010) and mites (Miller et al., 2013; Skoracka, Kuczynski, Szydlo, & Rector, 2013), some of which are characterised by cryptic species complexes. Discovering cryptic diversity has important consequences for estimates of global biodiversity, conservation planning, and the management of pests and diseases. Morphologically similar species can vary in pest status due to differences in genotypic and/or phenotypic traits that influence their host range and specificity, geographic distribution, the ability to vector diseases, or insecticide resistance (Ashfaq et al., 2014; Miller et al., 2013; Umina, Hoffmann, & Weeks, 2004). Therefore, recognising cryptic species and the differences in their biology and ecology are essential for effective management, with important implications for public health, agriculture and trade.

The diamondback moth, *Plutella xylostella*, is the major pest of brassica crops worldwide, costing an estimated US$4 to US$5 billion annually in direct losses and management costs (Furlong, Wright, & Dosdall, 2013; Zalucki et al., 2012). Insecticide resistance is widespread in populations around the world, fuelling wide-ranging research to develop alternative management tactics (Furlong et al., 2013; Li, Feng, Liu, You, & Furlong, 2016). *Plutella xylostella* was initially recorded in Australia in the late 1800s and rapidly became a widespread pest of brassica vegetables, and then canola following its expanded production from the 1990s (Endersby, McKechnie, Ridland, & Weeks, 2006; Furlong et al., 2008). Recently, Landry and Hebert (2013) through mtDNA barcoding identified a cryptic lineage of *Plutella* in Australia not detected in previous molecular studies of *P. xylostella* (Delgado & Cook, 2009; Endersby et al., 2006, 2011; Pichon et al., 2006; Roux et al., 2007; Saw, Endersby, & McKechnie, 2006). Although external morphology was indistinguishable from *P. xylostella*, deep mtDNA divergence (8.6%), differences in genital morphology and endemism in Australia led them to describe a new species, *Plutella australiana* Landry & Hebert. *Plutella australiana* was originally collected together with *P. xylostella* in light trap samples in eastern Australia, suggesting at least some ecological overlap (Landry & Hebert, 2013), but its biology, ecology and pest status were unknown.

The management of *P. xylostella* in Australian brassica crops has been a significant challenge for decades (Baker, 2011; Furlong et al., 2008), but the discovery of *P. australiana* has made the relative abundance and pest status of both species in these crops uncertain. With rare exception, *P. xylostella* and allied species feed on plants in the order Brassicales, mainly within the family Brassicaceae, (Landry & Hebert, 2013; Robinson & Sattler, 2001; Sarfraz, Dosdall, & Keddie, 2006), implying that the host range of *P. australiana* may include cultivated brassicas. Widespread resistance to pyrethroid and organophosphate insecticides has been attributed to Australian populations of *P. xylostella* from all vegetable and canola production regions, which has led to ineffective control during outbreaks (Baker, 2011; Endersby, Ridland, & Hoffmann, 2008). *Plutella xylostella* is well known as a highly migratory insect with a high capacity for gene flow (Furlong et al., 2013; Li et al., 2016), facilitating the rapid spread of resistance alleles. Australian *P. xylostella* are thought to frequently disperse, based on indirect evidence from ecological and genetic studies (Endersby et al., 2006; Furlong et al., 2008; Ridland & Endersby, 2008). Most studies have found a lack of genetic variation across microsatellite loci and mitochondrial markers among Australian and New Zealand populations of *P. xylostella*, consistent with high gene flow and/or recent ancestry (Delgado & Cook, 2009; Endersby et al., 2006; Furlong et al., 2008; Saw et al., 2006). While species identification was not in question in these studies, somewhat inconsistent findings in two studies from eastern Australia using allozymes or SSR markers (Pichon et al., 2006; Roux et al., 2007) might reflect the confounding presence of *P. australiana* samples (Landry & Hebert, 2013). With these considerations, future management of *P. xylostella* in Australian crops will require thorough understanding of the ecological requirements, genetic traits and pest status of the two *Plutella* species. Further, reproductive isolation between species is unknown but has implications for evolutionary inference and the potential for gene flow. The capacity for hybridization and introgression can lead to the exchange of insecticide resistance or other adaptive alleles (Clarkson et al., 2014; Hedrick, 2013).

Although mtDNA markers are widely used in studies of species identity and population structure (Ashfaq & Hebert, 2016; Hebert, Penton, Burns, Janzen, & Hallwachs, 2004; Smith et al., 2012), mitochondrial variation within or between species can be influenced by direct and/or indirect selection, or introgressive hybridization (Dupont, Porco, Symondson, & Roy, 2016; Whitworth, Dawson, Magalon, & Baudry, 2007). Interpretations of population history based on mtDNA can be corroborated using independent nuclear markers and/or integrative approaches (Roe & Sperling, 2007). One factor that can confound mtDNA is interactions with inherited bacterial symbionts (Hurst & Jiggins, 2005; Ritter et al., 2013). *Wolbachia* is an extremely widespread endosymbiont thought to infect at least half of arthropod (Weinert, Araujo-Jnr, Ahmed, & Welch, 2015) and 80% of lepidoptera (Ahmed, Breinholt, & Kawahara, 2016) species. It is mainly transmitted vertically from infected females to their offspring through the egg cytoplasm, and inheritance is therefore linked with mtDNA. To facilitate its spread, *Wolbachia* manipulates host reproductive biology to favour the fitness of infected females, by inducing host phenotypes that distort sex ratios (through feminization of males, male-killing or induction of parthenogenesis) or cause sperm-egg cytoplasmic incompatibility (CI) (Engelstaedter & Hurst, 2009; Werren, Baldo, & Clark, 2008). In the simple case involving a single CI-inducing strain, crosses with infected females are fertile but crosses between uninfected females and infected males fail to produce offspring. If maternal transmission is efficient and infected females have a reproductive advantage, *Wolbachia* infection can rapidly spread through insect populations (F. M. Jiggins, 2017), driving a selective sweep of a single haplotype and reducing mtDNA diversity (Shoemaker, Dyer, Ahrens, McAbee, & Jaenike, 2004). Very limited surveying to date has identified *Wolbachia* strains infecting *P. xylostella* at low frequency in populations from North America, Africa, Asia and Europe (Batista, Keddie, Dosdall, & Harris, 2010; Delgado & Cook, 2009; Jeyaprakash & Hoy, 2000). Because symbionts can contribute to reproductive isolation and shape mtDNA diversity (Hurst & Jiggins, 2005; Telschow, Hilgenboecker, Hammerstein, & Werren, 2014), assessing their role can provide important insights into host evolution and population structure (Dumas et al., 2013; Munoz, Baxter, Linares, & Jiggins, 2011; Ritter et al., 2013; X.-J. Sun, Xiao, Cook, Feng, & Huang, 2011).

Here we investigated the biology, ecology and genetic structure of these cryptic populations by collecting *Plutella* from brassicaceous plants throughout Australia and screening individuals to identify mtDNA lineages and *Wolbachia* infections. For a subset of populations, we examined genetic diversity using thousands of nuclear SNPs from across the genome. In addition, we assessed reproductive compatibility in laboratory crosses and determined the susceptibility of each species to commercial insecticides.

## Materials and methods

### Sample collection

*Plutella* was collected from canola crops, *Brassica* vegetable crops, forage brassicas and wild brassicas throughout Australia between March 2014 and December 2015. The wild species included wild radish, *Raphanus raphanistrum*, turnip weed, *Rapistrum rugosum*, sea rocket, *Cakile maritima*, Ward’s weed, *Carrichtera annua* and mixed stands of sand rocket, *Diplotaxis tenuifolia* and wall rocket, *D. muralis*. At each location, a sample of at least 25 individual larvae (rarely, eggs or pupae) were collected from randomly selected plants to achieve a representative sample. Samples were collected from Brassica vegetables by hand, from sea rocket by beating plants over a collection tray, from other hosts using a sweep net. Each population sample was separately reared in ventilated plastic containers on leaves of the original host material for 1-2 days and thereafter on cabbage leaves. Non-parasitised pupae or late-instar larvae were fresh frozen at — 80 °C. Pupae were visually sexed under a dissecting microscope.

### DNA isolation and COI genotyping

Individual pupae (but not larvae) were sexed under a dissecting microscope, then genomic DNA was isolated by homogenising whole individuals followed by two phenol and one chloroform extractions according to Zraket, Barth, Heckel, and Abbott (1990). DNA was treated with RNase A, then precipitated and re-suspended in TE buffer. *Plutella* lineages were distinguished using a PCR-RFLP assay (Perry, Pederson, & Baxter, 2017). A 707 bp COI region was amplified using a combination of two primer pairs: (i) PxCOIF (5′-TCAACAAATCATAAAGATATT-GG-3′) and PxCOIR (5′-TAAACTTCAGGGTGACCAAAAAATCA-3′), and (ii) PaCOIF (5^;^-TCAACAAATCATAAGGATATTGG-3′) and PaCOIR (5′-TAAACCTCTGGATGGCCAAA-AAATCA-3′). Ten microliter reactions were run with 2uL of MyTaq 5x buffer, 0.2uL of each primer (10 mm stocks), 1uL of DNA (approx. 5 ng) and 0.05 uL of MyTaq polymerase (Bioline). Samples were amplified at 95 °C for 2 minutes, then 35 cycles at 95 °C for 10 seconds, 52 °C for 20 seconds, 72 °C for 30 seconds followed by a 5 minute final extension at 72 °C. PCR products were digested at 37 °C for 1 hour with 1 unit of *AccI* (NEB) restriction enzyme with 2uL Cutsmart Buffer in a 20 uL reaction. Following digestion, products were separated using agarose gel electrophoresis (1.5%). *Plutella xylostella* products are approximately 516 bp and 191 bp and *P. australiana* products are 348 bp and 359 bp. COI amplicon sequencing was performed at the Australian Genome Research Facility (AGRF). In addition, we downloaded and re-analysed sequence trace files from Landry and Hebert (2013) (<u>dx.doi.org/10.5883/DS-PLUT1</u>) in geneious v10.0.6 (Kearse et al., 2012). Haplotype networks were constructed using R package pegas v0.9 (Paradis, 2010).

### *Wolbachia* screening and phylogenetics

*Wolbachia* infection was detected using two separate PCR assays of the 16S rRNA gene (16S-2 and 16S-6) according to Simoes, Mialdea, Reiss, Sagot, and Charlat (2011). To identify *Wolbachia* strains, the *Wolbachia surface protein (wsp)* gene was sequenced for a subset of individuals. Amplification was performed using wsp81F and wsp691R sequence primers (Zhou, Rousset, & O’Neill, 1998). Amplicons were sequenced using the reverse primer and aligned in geneious v10.0.6 (Kearse et al., 2012). We used a 493 bp alignment to construct a maximum likelihood phylogeny in RAxML v8.2.4 (Stamatakis, 2014) using a general time reversal substitution model with 1000 bootstraps.

### RADseq library preparation and sequencing

Libraries were prepared for restriction-site-associated DNA sequencing (RADseq) according to a protocol modified from Baird et al. (2008). Genomic DNA was quantified using a Qubit 2.0 fluorometer (Invitrogen) and 200 ng digested with 10 units of high fidelity *SbfI* in Cutsmart buffer (NEB) for 1 hour at 37 °C, then heat inactivated at 80 °C for 20 minutes. One microlitre of P1 adapter (100 nM) with a 6-base molecular identifier (MID) (top strand 5′-TCGTCGGCAGCG-TCAGATGTGTATAAGAGACAGxxxxxxtgca-3′, bottom strand 5′-[P]xxxxxxCTGTCTCTT-ATACACATCTGACGCTGCCGACGA-3′, x represents sites for MIDs) were then added using 0.5 uL T4 DNA ligase (Promega), 1nM ATP and Cutsmart buffer. Library pools were sheared using a Bioruptor sonicator (Diagenode), ends repaired (NEB), adenine overhangs added then P2 adapters (top strand 5′-[P]CTGTCTCTTATACACATCTCCAGAATAG-3′, bottom strand 5′-GTCTCGTGGGCTCGGAGATGTGTATAAGAGACAGT-3′) ligated. DNA purification between steps was performed using a MinElute PCR purification kit (Qiagen). Libraries were amplified using KAPA HiFi Hotstart Readymix (Kapa Biosystems) and Nextera i7 and i5 indexed primers with PCR conditions: 95 °C for 3 minutes, two cycles of 98 °C for 20 seconds, 54 °C for 15 seconds, 72 °C for 1 minute, then 15 cycles of 98 °C for 20 seconds, 65 °C for 15 seconds, 72 °C for 1 minute followed by a final extension of 72 °C for 5 minutes. Libraries were size-selected (300-700 bp) on agarose gel and purified using a minElute Gel Extraction Kit (Qiagen), then Illumina paired-end sequencing performed using HiSeq2500 (100 bp) or NextSeq500 (75 bp) at the AGRF.

### Read filtering and variant calling

Sequence reads were demultiplexed using radtools vl.2.4 (Baxter et al., 2011) allowing one base MID mismatch, then trimmomatic v0.32 (Bolger, Lohse, & Usadel, 2014) used to remove restriction sites, adapter sequences, a thymine base from reverse reads introduced by the P2 adapter, and quality filter using the illuminaclip tool with parameters: trailing:10 slid-ingwindow:4:15 minlen:40. Paired reads were aligned to the *P. xylostella* reference genome (accession number: GCF 000330985.1) using stampy v1.0.21 (Lunter & Goodson, 2011) with-baq and -gatkcigarworkaround options and expected substitution rate set to 0.03 for *P. xylostella* and 0.05 for *P. australiana*. Duplicate reads were removed using picard v1.71 (http://broadinstitute.github.io/picard/). Genotypes were called using the genome analysis tool kit v3.3-0 (DePristo et al., 2011; McKenna et al., 2010) haplotypecaller tool. We determined that base quality score recalibration using bootstrapped SNP databases was inappropriate for this dataset as it globally reduced quality scores. For downstream comparisons between species, we joint-genotyped *P. australiana* and *P. xylostella* individuals using the geno-typeGVCFs workflow. To examine finer scale population structure, we also joint-genotyped the *P. australiana* individuals alone. All variant callsets were hard-filtered with identical parameters using vcftools v0.1.12a Danecek et al. (2011): We removed indels and retained confidently called biallelic SNPs (GQ≤30) genotyped in at least 70% of individuals with a minimum genotype depth of 5, minQ≤400, average site depth of 12–100, minimum minor allele frequency of 0.05, in Hardy-Weinberg equilibrium at alpha = 0.05, retaining SNPs separated by a minimum of 2000 bp using the vcftools -thin function. To estimate genetic diversity, we generated a set of confidently called (GQ≤30) variant and invariant sites, and hard filtered to remove sites within repetitive regions and retain sites genotyped in at least 70 % of individuals with average site depth of 12–100.

### Genetic diversity and population structure

Genetic diversity was calculated for *Plutella* populations of both species from five locations. The R package hierfstat (Goudet & Jombart, 2015) was used to calculate observed heterozygosity, gene diversity and the inbreeding coefficient, F_is_, according to Nei (1987). Population means for site depth and number of SNPs, indels and private sites were calculated using the –depth function and vcfstats module in vcftools v0.1.12a (Danecek et al., 2011). Levels of heterozy-gosity sites within individuals were determined from all confidently called sites excluding indels using a custom python script parseVCF.py (https://github.com/simonhmartin) and visualised using R (R Core Team, 2017).

To examine population structure in *P. australiana*, a global estimate of F_st_ (Weir & Cock-erham, 1984) with bootstrapped 99% confidence intervals (10^4^ bootstraps) was calculated in R package diveRsity (Keenan, McGinnity, Cross, Crozier, & Prodoehl, 2013). Pairwise F_st_ values for all population pairs were calculated and significance determined using exact G tests (10^4^ mc burnins, 10^3^ batches, and 10^4^ iterations per batch) in genepop v4.6 (Rousset, 2008) after bonferroni correction for multiple comparisons.

Separate analysis of population structure was performed using the Bayesian clustering program structure v2.3.4 (Pritchard, Stephens, & Donnelly, 2000), first for all individuals of co-occurring *Plutella* species, and second for *P. australiana* alone. For all runs, we used 5 x 10^5^ burnins and 10^6^ MCMC replicates and performed ten independent runs for each K value from 1 to 10, where K is the number of genotypic clusters, using a different random seed for each run, assuming the locprior model with correlated allele frequencies and λ set to 1. The optimal value of K was determined using the deltaK method (Evanno, Regnaut, & Goudet, 2005) implemented in structure harvester (Earl & vonHoldt, 2012) and inspection of the likelihood distribution for each model. Q-matrices were aligned across runs using clumpp v1.1.2 (Jakobsson & Rosenberg, 2007) and visualised using distruct v1.1 (Rosenberg, 2004).

### Laboratory cultures of *Plutella* species

Laboratory cultures of *P. australiana* and *P. xylostella* were established from field populations and used for crossing experiments and insecticide bioassays. *Plutella* adults were collected at light traps at Angle Vale and Urrbrae, South Australia, in October–November 2015. Females were isolated and allowed to lay eggs, then identified using PCR-RFLP and progeny pooled to produce separate cultures of each species. A laboratory culture of the Waite Susceptible *P. xylostella* strain (S) has been maintained on cabbage without insecticide exposure for approximately 24 years (≈ 305 generations). All cultures were maintained in laboratory cages at 26 ± 2.0 °C and a 14:10 (L:D) hour photoperiod at the South Australian Research and Development Institute, Waite Campus, Adelaide, South Australia. The *P. australiana* culture was maintained on sand rocket, *Diplotaxis tenuifolia* L., and the *P. xylostella* culture maintained on cabbage, *Brassica oleracea* L. var. *capitata*. The purity of cultures was assessed regularly using PCR-RFLP.

### Crossing experiments

*Plutella australiana* and *P. xylostella* pupae were sexed under a stereo microscope, then placed into individual 5 mL clear polystyrene tubes with fine mesh lids and gender visually confirmed after eclosion. Enclosures used for crossing experiments were 850mL polypropylene pots (Bon-son Pty Ltd) modified with lateral holes covered with voile mesh for ventilation. Crosses of single mating pairs were performed on laboratory benches at 26 ± 2.0 °C and 14:10 (L:D) photoperiod using 3-week old *D. tenuifolia* seedlings as the host plant. After seven days, adults were collected into a 1.5 mL tube and fresh frozen at -80 °C for species confirmation using PCR-RFLP. Seedlings were examined and eggs counted under a stereo microscope, then returned to enclosures to allow egg hatch. Larvae were provided with fresh 3–4 week old seedlings until pupation, then pupae were individually collected into 5mL tubes. Hybrid F1xF1 crosses and back-crosses were then performed as above. The presence of egg and adult offspring was recorded for all replicates, and for the majority of replicates (> 80%), the numbers of offspring were counted and used to calculate a mean.

### Insecticide bioassays

Insecticide susceptibility of field-collected *Plutella* strains was compared to the susceptible *P. xylostella* (S) reference in dose-response assays using four commercial insecticides: Dominex (100 g L^-1^ alpha-cypermethrin), Proclaim (44 g kg^-1^ emamectin benzoate), Coragen (200 g L^-1^ chlorantraniliprole) and Success Neo (120 g L^-1^ spinetoram). Bioassays were performed by placing 3^rd^ instar larvae onto inverted leaf discs embedded in 1% agar in 90mm Petri dishes. Cabbage leaves, *Brassica oleracea* L. var. *capitata* were used for *P. xylostella* and canola leaves, *B. napus* L. var. ‘ATR Stingray’ were used for *P. australiana*. Eight concentrations and a water-only control were evaluated for each insecticide using four replicates of ten larvae. A 4 mL aliquot of test solution was applied directly to leaves using a Potter Spray Tower (Burkard Manufacturing Co. Ltd.) calibrated to deliver an aliquot of 3.52 ± 0.09 mg cm^-2^. After application, each dish was placed in a controlled temperature room at 25 ± 0.5 °C, then mortality assessed after 48 hours (Dominex, Success Neo) or 72 hours (Proclaim, Coragen). Dose-response analysis was performed using log-logistic regression in R package drc (Ritz, Baty, Streibig, & Gerhard, 2015), and the fitted models were used to estimate the lethal concentration predicted to cause 50% (LC_50_) and 99 % (LC_99_) mortality of the test population with 95%. Resistance ratios were calculated by dividing the LC_50_ and LC_99_ estimates for field strains by the corresponding LC estimates for the *P. xylostella* (S) reference. In addition, we calculated a ratio of the commercial field concentration with the LC_99_ estimates, based on the field application rates of each insecticide registered for use against *P. xylostella* in Australian brassica vegetable crops: 40 mg ha^-1^ a.i. (Dominex), 13.2 mg ha^-1^ a.i. (Proclaim), 20 mg ha^-1^ a.i. (Coragen) and 48mg ha^-1^ a.i. (Success Neo).

## Results

### Geographic distribution and host association of *Plutella* species

*Plutella* larvae were collected from brassica plants at 75 locations in Australia and 1477 individuals were genotyped at the COI locus using PCR-RFLP to identify species. Of these, 88% (n=1300) were genotyped as *P. xylostella*, 10% (n=147) as *P. australiana*, and 2% (n=30) were unresolved (Table 1). *Plutella australiana* was identified in around one quarter (n=20/75) of collections distributed across southern Australia, while *P. xylostella* occurred at all locations except Cunnamulla, Queensland, in a collection from wild African mustard, Brassica tournefortii (Table 1). The sex ratio was not different from 1:1 for *P. xylostella* (481 females, 517 males, χ^2^=1.2986, p=0.2545) or *P. australiana* (63 females, 55 males, χ^2^=0.5424, p=0.4615). The relative incidence and abundance of *P. australiana* was >2-fold higher in the eastern state of New South Wales than in other states (Table 2, Figure 1). *Plutella australiana* larvae were detected in 29% (n=5/17) of collections from wild species, including wild radish, Raphanus raphanistrum, wild turnip, Rapistrum rugosum and mixed stands of sand rocket, *D. tenuifolia* and wall rocket, D. muralis. Among cultivated crops, *P. australiana* larvae occurred in 36% (n=14/39) of samples from canola, consisting of 11% of total *Plutella* individuals, but were not identified from commercial brassica vegetable farms (Table 2). However, *P. australiana* eggs were collected from kale on one farm.

**Figure 1:**
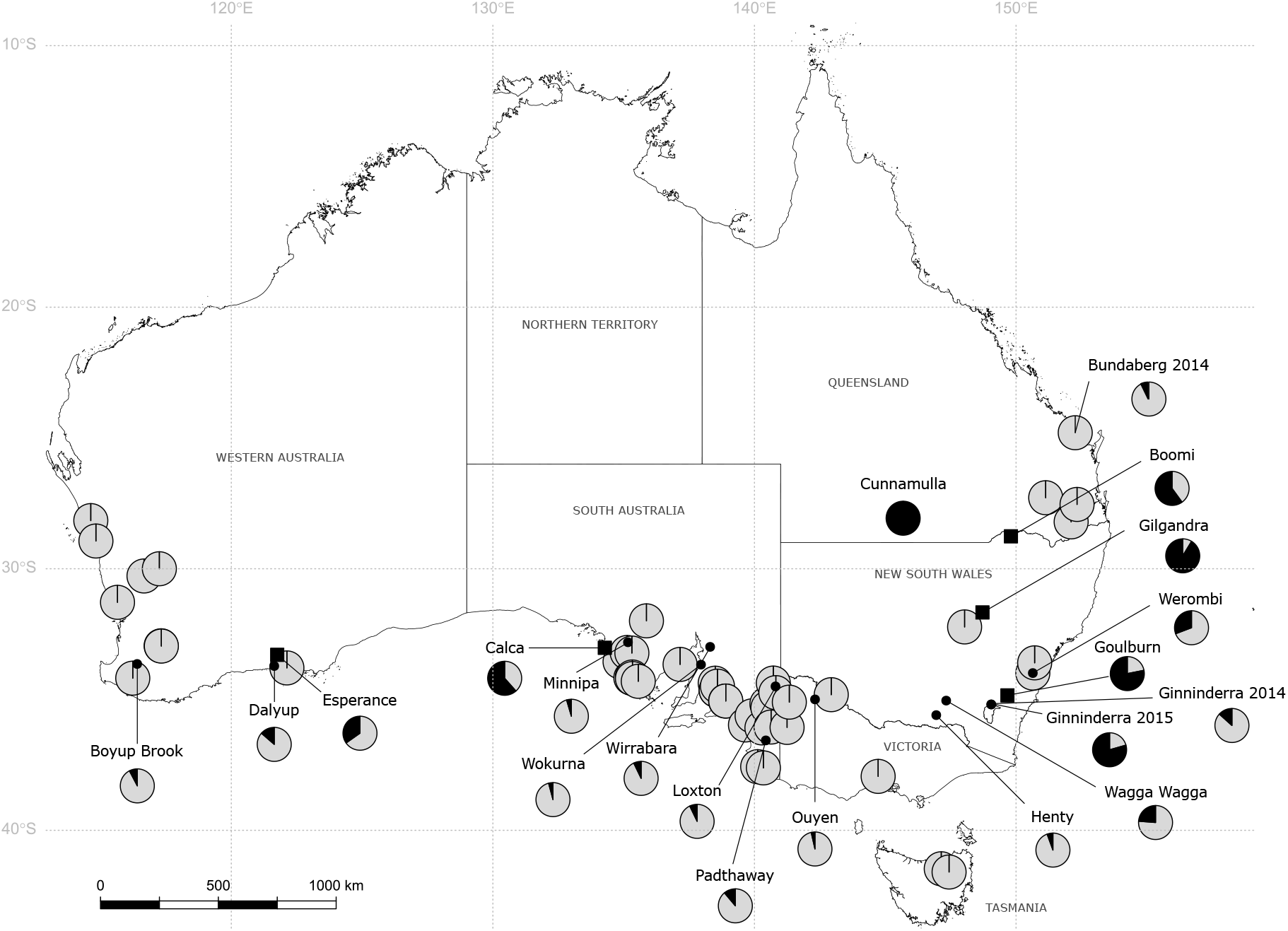
The distribution of *P. xylostella* (light grey) and *P. australiana* individuals (black) in larval collections from brassicaceous plants in Australia during 2014 and 2015. Pie diagrams show the relative proportion of each species at each location. Overlapped pies represent locations with 100 % *P. xylostella*. Black squares indicate five locations where individuals of each species were RAD sequenced.

**Table 1:** Summary of *Plutella* collections from Australia. For each location, the numbers and frequency (*f*) of each species and the *Wolbachia* infection status of *P. xylostella* are presented. All *P. australiana* individuals were infected with *Wolbachia*.

| Location | Collection date | Latitude | Longitude | Host | No. genotyped | *P. australiana* | *P. xylostella* |  |
| --- | --- | --- | --- | --- | --- | --- | --- | --- |
|  |  |  |  |  |  | No. (*f*) | No. (*f*) | No. (*f*)<br>*wol*-infected |
| Boomi NSW | Sep-14 | −28.76° | 149.81° | Canola | 25 | 15 (0.60) | 10 (0.40) | 0 (0.00) |
| Gilgandra NSW | Sep-14 | −31.67° | 148.72° | Wild turnip | 23 | 21 (0.91) | 2 (0.09) | 0 (0.00) |
| Ginninderra NSW | Oct-15 | −35.19° | 149.05° | Canola | 34 | 27 (0.79) | 7 (0.21) | 0 (0.00) |
| Ginninderra NSW | Sep-14 | −35.19° | 149.05° | Canola | 15 | 2 (0.13) | 13 (0.87) | 0 (0.00) |
| Goulburn NSW | Nov-15 | −34.84° | 149.67° | Canola | 32 | 25 (0.78) | 7 (0.22) | 0 (0.00) |
| Henty NSW | Oct-14 | −35.60° | 146.95° | Canola | 18 | 1 (0.06) | 17 (0.94) | 0 (0.00) |
| Narromine NSW | Sep-14 | −32.22° | 148.03° | Canola | 26 | 0 (0.00) | 26 (1.00) | 1 (0.04) |
| Richmond NSW | Oct-15 | −33.60° | 150.71° | Cabbage | 21 | 0 (0.00) | 21 (1.00) | 0 (0.00) |
| Wagga Wagga NSW | Sep-14 | −35.04° | 147.33° | Canola | 21 | 5 (0.24) | 16 (0.76) | 0 (0.00) |
| Werombi NSW | Oct-15 | −34.00° | 150.56° | Kale | 13 | 4 (0.31) | 9 (0.69) | 0 (0.00) |
| Werombi NSW | Nov-14 | −33.99° | 150.64° | Vegetables | 16 | 0 (0.00) | 16 (1.00) | 0 (0.00) |
| Bundaberg QLD | Oct-14 | −24.80° | 152.26° | Canola | 14 | 1 (0.07) | 13 (0.93) | 0 (0.00) |
| Bundaberg QLD | Sep-15 | −24.80° | 152.26° | Canola | 30 | 0 (0.00) | 30 (1.00) | 0 (0.00) |
| Cunnamulla QLD | Sep-15 | −28.07° | 145.68° | African mustard | 17 | 17 (1.00) | 0 (0.00) | 0 |
| Dalby QLD | Sep-14 | −27.28° | 151.13° | Canola | 30 | 0 (0.00) | 30 (1.00) | 0 (0.00) |
| Gatton QLD | Oct-14 | −27.54° | 152.33° | Broccoli | 16 | 0 (0.00) | 16 (1.00) | 0 (0.00) |
| Gatton QLD | Nov-15 | −27.54° | 152.33° | Broccoli | 15 | 0 (0.00) | 15 (1.00) | 0 (0.00) |
| Warwick QLD | Oct-15 | −28.21° | 152.11° | Canola | 16 | 0 (0.00) | 16 (1.00) | 0 (0.00) |
| Calca SA | Apr-14 | −33.02° | 134.28° | Sand rocket, Wall rocket | 13 | 8 (0.62) | 5 (0.38) | 0 (0.00) |
| Cocata SA | Sep-14 | −33.20° | 135.13° | Canola | 18 | 0 (0.00) | 18 (1.00) | 0 (0.00) |
| Colebatch SA | Feb-15 | −35.97° | 139.66° | Forage brassica | 18 | 0 (0.00) | 18 (1.00) | 0 (0.00) |
| Coonalpyn SA | Oct-15 | −35.62° | 139.91° | Wild radish | 11 | 0 (0.00) | 11 (1.00) | 0 (0.00) |
| Cowell SA | Sep-14 | −33.66° | 137.16° | Canola | 16 | 0 (0.00) | 16 (1.00) | 0 (0.00) |
| Keith SA | Oct-14 | −36.09° | 140.29° | Canola | 32 | 0 (0.00) | 32 (1.00) | 6 (0.19) |
| Lameroo SA | Sep-14 | −35.32° | 140.51° | Canola | 16 | 0 (0.00) | 16 (1.00) | 0 (0.00) |
| Lameroo SA | Oct-15 | −35.17° | 140.48° | Canola | 14 | 0 (0.00) | 14 (1.00) | 0 (0.00) |
| Littlehampton SA | Oct-14 | −35.06° | 138.90° | Cabbage | 34 | 0 (0.00) | 34 (1.00) | 6 (0.18) |
| Littlehampton SA | Sep-15 | −35.06° | 138.90° | Brussels sprouts | 8 | 0 (0.00) | 8 (1.00) | 0 (0.00) |
| Loxton SA | Oct-15 | −34.50° | 140.80° | Canola | 14 | 1 (0.07) | 13 (0.93) | 0 (0.00) |
| Loxton SA | Sep-14 | −34.37° | 140.72° | Canola | 31 | 0 (0.00) | 31 (1.00) | 0 (0.00) |
| Mallala SA | Sep-15 | −34.38° | 138.50° | Canola | 26 | 0 (0.00) | 26 (1.00) | 0 (0.00) |
| Meribah SA | Sep-14 | −34.74° | 140.82° | Canola | 16 | 0 (0.00) | 16 (1.00) | 0 (0.00) |
| Millicent SA | Apr-15 | −37.61° | 140.34° | Canola | 9 | 0 (0.00) | 9 (1.00) | 2 (0.22) |
| Minnipa SA | Oct-15 | −32.81° | 135.16° | Canola | 22 | 1 (0.05) | 21 (0.95) | 0 (0.00) |
| Moonaree SA | Aug-14 | −31.99° | 135.87° | Ward's weed | 16 | 0 (0.00) | 16 (1.00) | 0 (0.00) |
| Mt Hope SA | Sep-14 | −34.14° | 135.33° | Canola | 29 | 0 (0.00) | 29 (1.00) | 0 (0.00) |
| Mt Hope SA | Sep-15 | −34.20° | 135.34° | Canola | 16 | 0 (0.00) | 16 (1.00) | 0 (0.00) |
| Padthaway SA | Oct-15 | −36.56° | 140.43° | Canola | 18 | 2 (0.11) | 16 (0.89) | 0 (0.00) |
| Picnic Beach SA | Sep-14 | −34.17° | 135.27° | Sea rocket | 16 | 0 (0.00) | 16 (1.00) | 0 (0.00) |
| Picnic Beach SA | Apr-14 | −34.17° | 135.27° | Sea rocket | 2 | 0 (0.00) | 2 (1.00) | 0 (0.00) |
| Redbanks SA | Oct-14 | −34.49° | 138.59° | Canola | 38 | 0 (0.00) | 38 (1.00) | 1 (0.03) |
| Sherwood SA | Oct-14 | −36.05° | 140.64° | Wild radish | 8 | 0 (0.00) | 8 (1.00) | 0 (0.00) |
| Southend SA | Apr-15 | −37.57° | 140.12° | Sea rocket | 18 | 0 (0.00) | 18 (1.00) | 0 (0.00) |
| Tintinara SA | Oct-15 | −35.97° | 139.66° | Forage brassica | 17 | 0 (0.00) | 17 (1.00) | 0 (0.00) |
| Ucontichie SA | Sep-14 | −33.22° | 135.31° | Canola | 3 | 0 (0.00) | 3 (1.00) | 0 (0.00) |
| Virginia SA | Oct-14 | −34.64° | 138.54° | Broccoli | 18 | 0 (0.00) | 18 (1.00) | 1 (0.06) |
| Virginia SA | Sep-15 | −34.64° | 138.54° | Cabbage | 23 | 0 (0.00) | 23 (1.00) | 0 (0.00) |
| Walkers Beach SA | Sep-14 | −33.55° | 134.86° | Sea rocket | 16 | 0 (0.00) | 16 (1.00) | 0 (0.00) |
| Walkers Beach SA | Mar-15 | −33.55° | 134.86° | Sea rocket | 16 | 0 (0.00) | 16 (1.00) | 0 (0.00) |
| Walkers Beach SA | Sep-15 | −33.55° | 134.86° | Sea rocket | 19 | 0 (0.00) | 19 (1.00) | 0 (0.00) |
| Wirrabara SA | Oct-14 | −32.99° | 138.31° | Canola | 28 | 2 (0.07) | 26 (0.93) | 0 (0.00) |
| Wokurna SA | Sep-15 | −33.67° | 137.96° | Wild radish | 24 | 1 (0.04) | 23 (0.96) | 0 (0.00) |
| Wurramunda SA | Apr-14 | −34.30° | 135.56° | Wild canola | 16 | 0 (0.00) | 16 (1.00) | 0 (0.00) |
| Deddington TAS | Nov-14 | −41.59° | 147.44° | Kale | 16 | 0 (0.00) | 16 (1.00) | 0 (0.00) |
| Launceston TAS | Nov-14 | −41.47° | 147.14° | Wild mustard | 16 | 0 (0.00) | 16 (1.00) | 0 (0.00) |
| Newstead TAS | Nov-15 | −41.59° | 147.44° | Cauliflower | 22 | 0 (0.00) | 22 (1.00) | 0 (0.00) |
| Cowangie VIC | Oct-15 | −35.10° | 141.33° | Canola | 19 | 0 (0.00) | 19 (1.00) | 0 (0.00) |
| Ouyen VIC | Sep-14 | −35.00° | 142.31° | Canola | 28 | 1 (0.04) | 27 (0.96) | 0 (0.00) |
| Robinvale VIC | Sep-14 | −34.81° | 142.94° | Canola | 16 | 0 (0.00) | 16 (1.00) | 0 (0.00) |
| Werribee VIC | Oct-14 | −37.94° | 144.73° | Cauliflower | 16 | 0 (0.00) | 16 (1.00) | 0 (0.00) |
| Werribee VIC | Nov-15 | −37.94° | 144.73° | Cauliflower | 16 | 0 (0.00) | 16 (1.00) | 0 (0.00) |
| Yanac VIC | Sep-14 | −36.06° | 141.25° | Canola | 17 | 0 (0.00) | 17 (1.00) | 0 (0.00) |
| Boyup Brook WA | Sep-14 | −33.64° | 116.40° | Canola | 26 | 2 (0.08) | 24 (0.92) | 0 (0.00) |
| Dalwallinu WA | Sep-15 | −30.28° | 116.66° | Canola | 20 | 0 (0.00) | 20 (1.00) | 0 (0.00) |
| Dalyup WA | Oct-15 | −33.72° | 121.64° | Wild radish | 22 | 3 (0.14) | 19 (0.86) | 0 (0.00) |
| Esperance WA | Sep-14 | −33.29° | 121.76° | Canola | 23 | 8 (0.35) | 15 (0.65) | 1 (0.07) |
| Esperance WA | Oct-15 | −33.79° | 122.13° | Canola | 16 | 0 (0.00) | 16 (1.00) | 0 (0.00) |
| Gingin WA | Dec-14 | −31.28° | 115.65° | Red cabbage | 23 | 0 (0.00) | 23 (1.00) | 1 (0.04) |
| Kalannie WA | Sep-15 | −30.00° | 117.25° | Canola | 18 | 0 (0.00) | 18 (1.00) | 0 (0.00) |
| Manjimup WA | Dec-14 | −34.18° | 116.23° | Chinese cabbage | 17 | 0 (0.00) | 17 (1.00) | 0 (0.00) |
| Manjimup WA | Nov-15 | −34.18° | 116.23° | Vegetables | 13 | 0 (0.00) | 13 (1.00) | 0 (0.00) |
| Narrogin WA | Oct-15 | −32.95° | 117.32° | Wild radish, Wild canola | 15 | 0 (0.00) | 15 (1.00) | 0 (0.00) |
| Narrogin WA | Oct-15 | −32.96° | 117.33° | Canola | 32 | 0 (0.00) | 32 (1.00) | 0 (0.00) |
| Walkaway WA | Sep-14 | −28.94° | 114.83° | Canola | 19 | 0 (0.00) | 19 (1.00) | 0 (0.00) |
| Walkaway WA | Sep-14 | −28.16° | 114.63° | Canola | 16 | 0 (0.00) | 16 (1.00) | 0 (0.00) |
| **Total** |  |  |  |  | **1447** | **147 (0.10)** | **1300 (0.90)** | **19 (0.01)** |

**Table 2:** Number and frequency (*f*) of *P. australiana* in *Plutella* collections from different Australian states and brassica host types.

| Group | No.<br>collection<br>locations | No. ($f$)<br>locations<br>*P. aus* | No.<br>individuals<br>genotyped | No. ($f$)<br>*P. aus* |
| --- | --- | --- | --- | --- |
| ***Australian state*** |  |  |  |  |
| New South Wales (NSW) | 11 | 8 (0.73) | 244 | 100 (0.41) |
| Queensland (QLD) | 7 | 2 (0.29) | 138 | 18 (0.13) |
| Western Australia (WA) | 13 | 3 (0.23) | 260 | 13 (0.05) |
| South Australia (SA) | 35 | 6 (0.17) | 639 | 15 (0.02) |
| Victoria (VIC) | 6 | 1 (0.17) | 112 | 1 (0.01) |
| Tasmania (TAS) | 3 | 0 (0.00) | 54 | 0 (0.00) |
| ***Brassica host type*** |  |  |  |  |
| Wild brassicas | 17 | 5 (0.29) | 268 | 50 (0.19) |
| Canola | 39 | 14 (0.36) | 848 | 93 (0.11) |
| Vegetables | 16 | 1 (0.06) | 287 | 4 (0.01) |
| forage | 3 | 0 (0.00) | 44 | 0 (0.00) |

### *Wolbachia* infections

*Plutella* individuals (n=1447) were screened for *Wolbachia* infection using 16S rRNA PCR assays. Only 1.5% (n=19/1300) of *P. xylostella* collected from eight different locations were infected (Table 1). In contrast, all 147 *P. australiana* individuals were infected with *Wolbachia* across the 20 locations where this species occurred. To identify Wolbachia strains, a Wolbachia surface protein (wsp) amplicon was sequenced from 14 *P. xylostella* and 30 *P. australiana* individuals. Each species was infected with a different strain. The wsp sequence for Australian *P. xylostella* showed 100 % identity to a *Wolbachia* supergroup A isolate infecting *P. xylostella* from Malaysia, plutWA1 (Delgado & Cook, 2009). For *P. australiana*, the wsp sequence showed 100 % identity to to a *Wolbachia* supergroup B isolate infecting a mosquito, Culex pipiens, from Turkey and the winter moth, Operophtera brumata, from the Netherlands (Figure 2).

**Figure 2:**
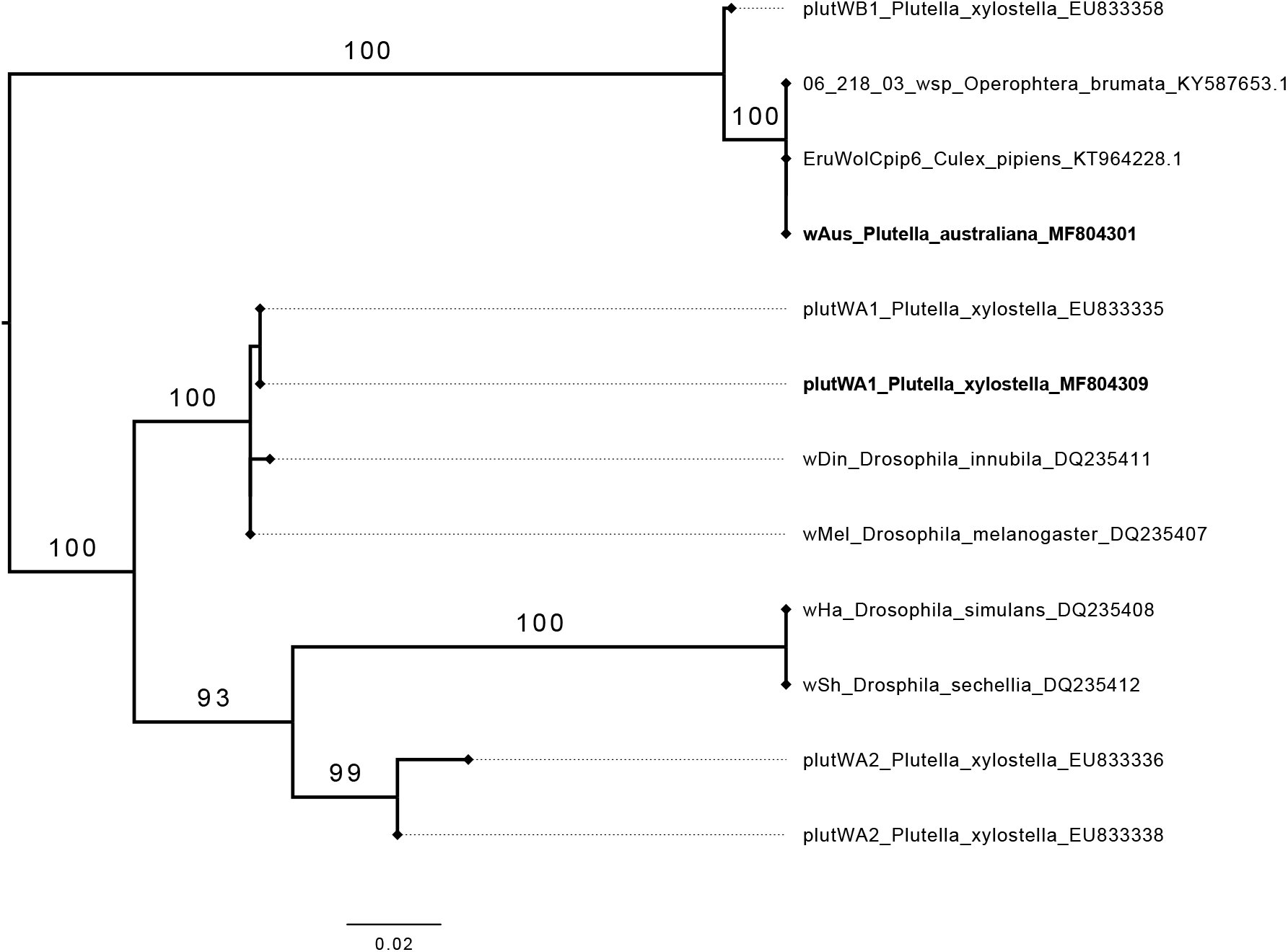
Maximum likelihood phylogeny of *Wolbachia wsp* amplicons for *Plutella* and other arthropods. The strain infecting *P. australiana* (*wAus*) was identical to a *Wolbachia* supergroup B strain reported from Culex pipiens and Operophtera brumata. The strain infecting Australian *P. xylostella* was identical to a supergroup A strain (*plutWA1*) reported from Malaysian *P. xylostella*. Labels include the *Wolbachia* strain, host species and GenBank accession number. Labels in bold denote strains sequenced in this study.

**Figure 3:**
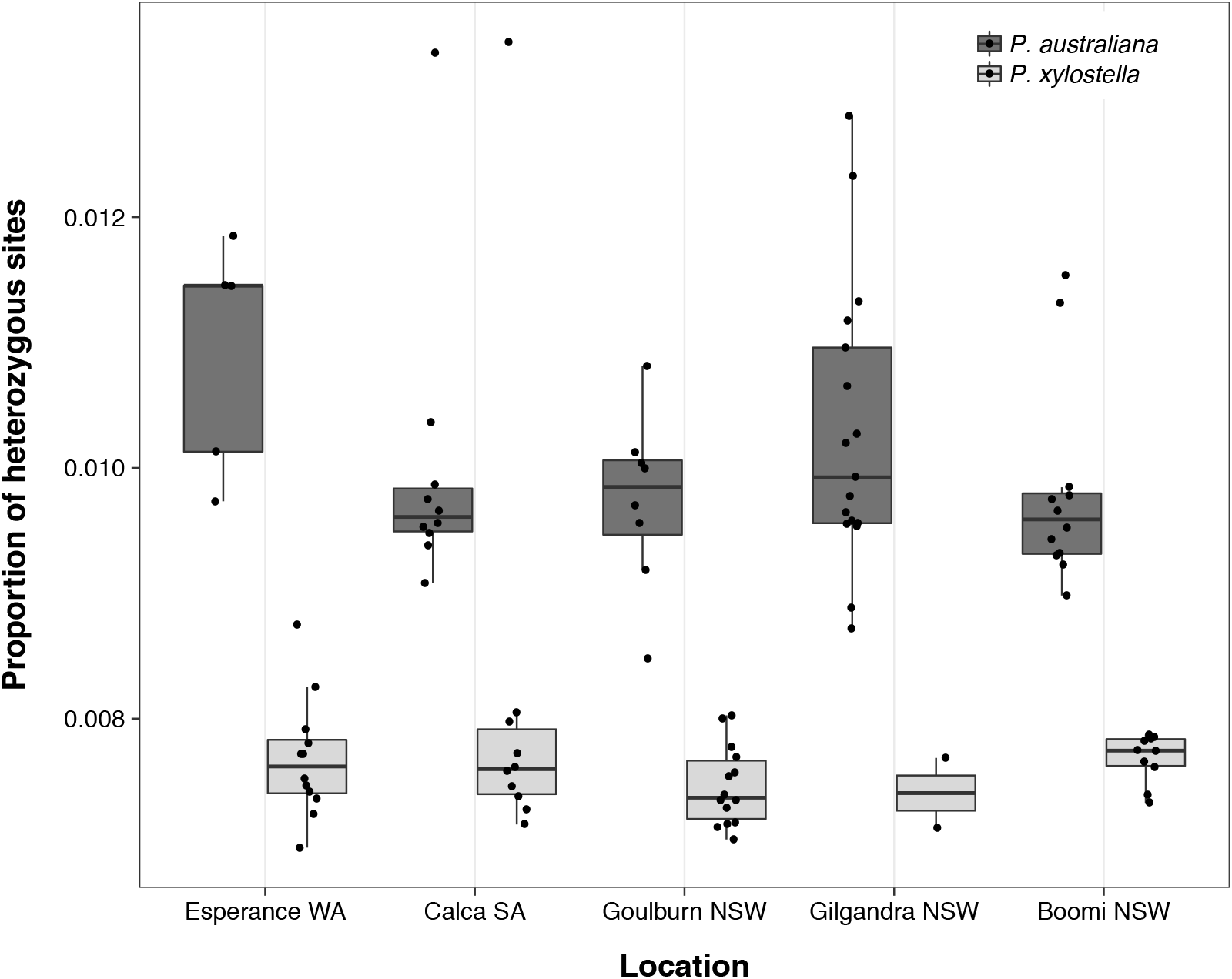
The proportion of heterozygous sites across 293 372 confidently called nuclear sites for individuals of *P. xylostella* (light grey boxes, n = 48) and *P. australiana* (dark grey boxes, n = 52) from five locations. Heterozygosity was consistently higher in *P. australiana*.

### Crossing experiments

Inter-species single pair mating experiments showed that hybridization between *P. australiana* and *P. xylostella* was possible, yet less successful than intra-species crosses. While most intra-species crosses produced adult offspring, the fecundity of *P. xylostella* was >2-fold higher than for *P. australiana* (Table 6). Both reciprocal inter-species crosses produced F1 adult offspring, but success was asymmetric and notably higher in the pairs with *P. australiana* females. In this direction, there was a strong male bias in the F1 progeny: from 76 cross replicates, 16 collectively produced 9 female and 80 male adults, a ratio of 8.9. Hybrid F1xF1 crosses for both parental lines produced F2 adult offspring. For the *P. australiana* maternal line, parental back-crosses using F1 hybrid males successfully produced offspring, while parental back-crosses with F1 hybrid females were sterile. For the *P. xylostella* maternal line, low fitness allowed only a single parental back-cross replicate, which involved a hybrid female and was sterile.

### Mitochondrial haplotype diversity

Mitochondrial haplotype networks of Australian *Plutella* were constructed using a 613 bp COI alignment that included 81 sequences from this study and 108 from Landry and Hebert (2013). We found low haplotype diversity within Australian *P. xylostella*, consistent with previous reports (Delgado & Cook, 2009; Juric, Salzburger, & Balmer, 2017; Saw et al., 2006). Only five haplotypes were identified among 102 individuals, including three identified by Saw et al. (2006) and three of which occurred in single individuals. The most common haplotype, PxCOI01, occurred at high frequency and differed by a single base from other haplotypes (Figure 5a, Table S1). Similarly, nine closely related haplotypes were identified in 87 *P. australiana* individuals, with seven occurring in single individuals (Figure 5B). The most common haplotype, PaCOI01, occurred at high frequency and differed by 1-2 bases from other haplotypes (Figure 5b, Table S2).

**Figure 4:**
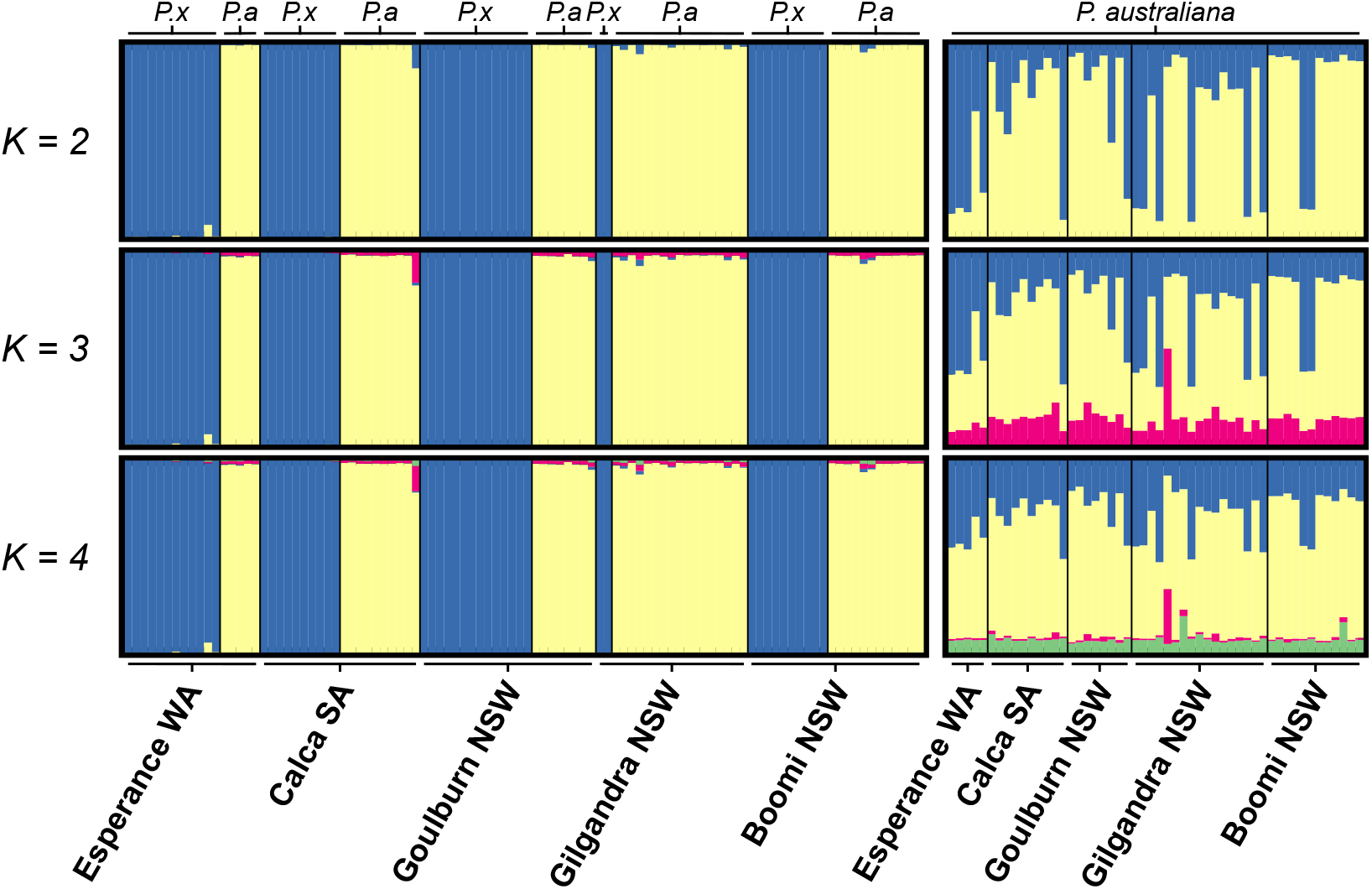
Proportional assignment of *Plutella* individuals to genotypic clusters, *K*, based on Bayesian structure analysis. Individuals are represented by vertical bars and genotypic clusters are represented by different colours. Bar plots are presented for *K* values from 2–4. Left panel: Analysis for 48 *P. xylostella* (labelled *P.x*) and 52 *P. australiana* (labelled *P.a*). Right panel: Analysis for 52 *P. australiana* individuals alone.

**Figure 5:**
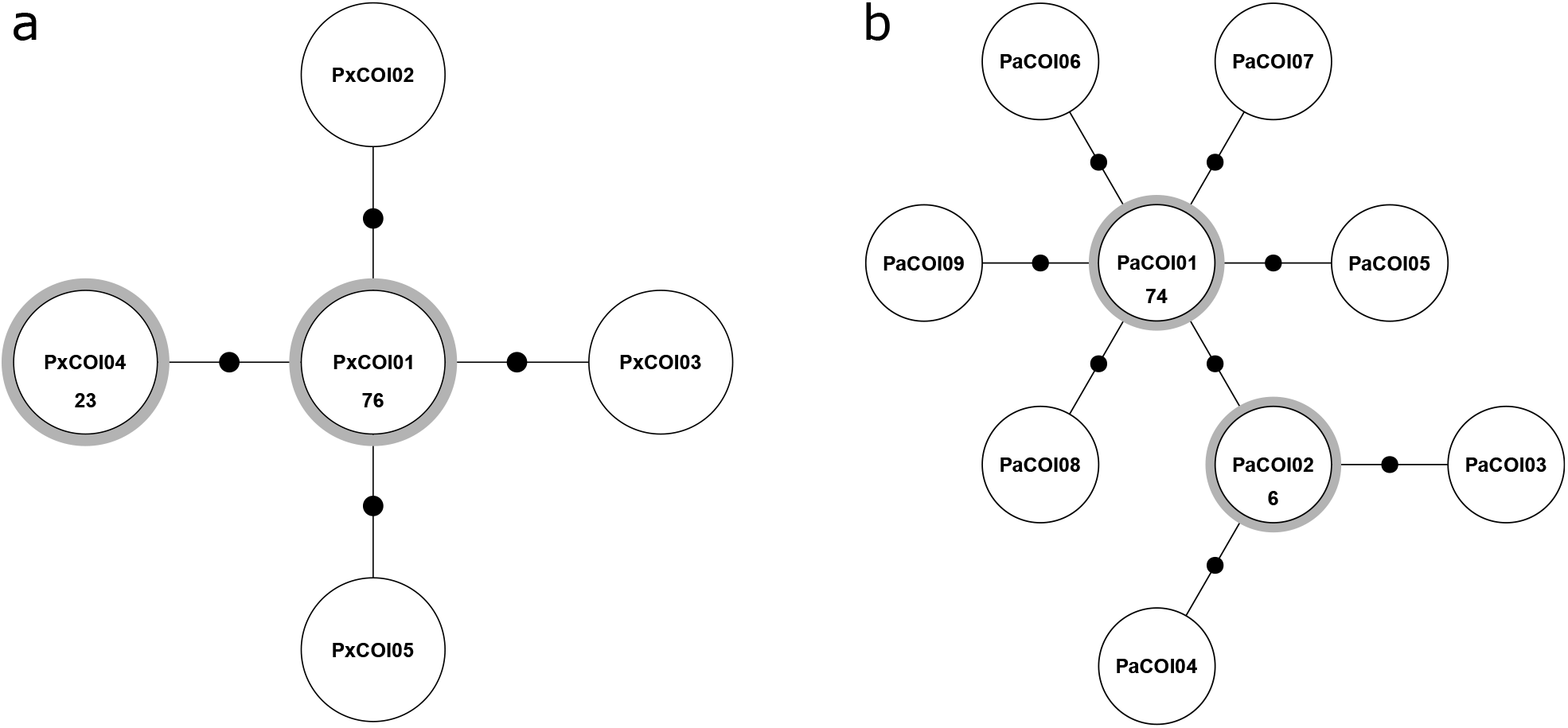
Haplotype network for *P. xylostella* (a) (n=102, this study 44) and *P. australiana* (b) (n=87, this study 37) individuals from Australia based on a 613 bp COI sequence alignment. Haplotypes shared by more than one individual are shown in circles with a grey border with the number of individuals indicated inside the circle. Small solid circles each denote one base difference between haplotypes

### Nuclear diversity and population structure

At five collection locations, *P. australiana* co-occurred with *P. xylostella* in sufficient numbers to enable comparison of nuclear genomes, though the relative abundance of species varied between locations. To ensure representation from the south-west region of Australia, the Esperance population (n=5) was formed by including one *P. australiana* individual from Boyup Brook. Despite only two *P. xylostella* individuals at Gilgandra, this population had 17 P. *australiana* individuals and was included. To generate nuclear markers, we performed RADseq for a total of 54 P. *australiana* and 48 P. *xylostella* individuals.

Illumina sequencing and demultiplexing using radtools (Baxter et al., 2011) yielded 282.7 million raw sequence reads. Two P. *australiana* individuals with low sequencing depth were excluded. Following read quality filtering and mapping, genotypes were called for 100 individuals from both species. Hard filtering retained 305 136 confidently called variant and invariant sites at a mean depth >36 per individual, and a subset of 707 widely-dispersed SNP variants (to avoid linkage bias), for comparative analyses of genetic diversity and population structure.

Analysis of nuclear diversity across 305 136 sites revealed a striking contrast between species, with notably higher diversity in populations of P. *australiana* than co-occurring populations of P. *xylostella* (Table 3). The mean observed heterozygosity within populations ranged from 0.13-0.16 for *P. australiana* and 0.009-0.010 for P. *xylostella*. Similarly, the average numbers of SNPs, indels and private alleles were considerably higher within *P. australiana* populations. As *P. australiana* may have fixed nucleotide differences relative to the P. *xylostella* reference genome that may affect population level statistics, we also removed indels from this dataset and directly compared the heterozygosity among individuals using 293 372 sites. This showed that *P. australiana* individuals had on average a > 1.5-fold higher proportion of heterozygous sites than P. *xylostella* individuals (Figure 3).

**Table 3:** Population statistics for variant and invariant sites for sympatric populations of *P. australiana* (*P. aus*) and *P. xylostella* (*P. x*) from five locations. Statistics presented include population means for the number of individuals genotyped per locus (*n*), observed heterozygosity (*H_o_*), gene diversity (*H_s_*) and Nei’s inbreeding coefficient, *F_IS_*.

| Population | Species | $n$ | Sites | Site<br>depth | SNPs | Indels | Private<br>sites | $H_o$ | $H_s$ | $F_{is}$ |
| --- | --- | --- | --- | --- | --- | --- | --- | --- | --- | --- |
| Boomi NSW | *P. aus* | 11.0 | 280 250 | 40 | 7356 | 1157 | 215 | 0.013 | 0.015 | 0.091 |
|  | *P. x* | 9.4 | 286 741 | 41 | 4431 | 568 | 30 | 0.009 | 0.010 | 0.041 |
| Calca SA | *P. aus* | 8.7 | 264 486 | 30 | 6771 | 1030 | 212 | 0.014 | 0.015 | 0.059 |
|  | *P. x* | 9.2 | 279 935 | 44 | 4354 | 576 | 43 | 0.010 | 0.011 | 0.009 |
| Esperance WA | *P. aus* | 4.5 | 272 426 | 28 | 6676 | 1037 | 214 | 0.016 | 0.015 | -0.032 |
|  | *P. x* | 11.0 | 278 874 | 35 | 4149 | 538 | 23 | 0.010 | 0.010 | 0.020 |
| Gilgandra NSW | *P. aus* | 15.6 | 280 533 | 39 | 7311 | 1131 | 216 | 0.014 | 0.015 | 0.080 |
|  | *P. x* | 1.9 | 281 509 | 42 | 4256 | 526 | 28 | 0.009 | 0.009 | -0.054 |
| Goulburn NSW | *P. aus* | 6.8 | 259 152 | 29 | 6607 | 1009 | 193 | 0.013 | 0.015 | 0.060 |
|  | *P. x* | 12.8 | 278 253 | 36 | 4156 | 530 | 26 | 0.009 | 0.010 | 0.053 |

Genetic structure was investigated using 707 nuclear SNPs for co-occurring populations of each species with the Bayesian clustering program structure. The deltaK method predicted a strong optimal at *K* = 2 genotypic clusters. *Plutella australiana* and P. *xylostella* individuals were clearly separated into distinct genotypic clusters in accordance to their mtDNA genotypes regardless of geographic location (Figure 4, left panel). A small degree of admixture can be seen for some individuals, as shown by sharing of coloured bars.

Assessing population structure from datasets with multiple species can mask heirachical structure (Kalinowski, 2011). To address this, genotypes were separately called for 52 P. australiana individuals, and hard filtering retained a set of 976 widely-dispersed SNP variants at a mean depth >32 per individual for examination of finer scale structure among five populations. The deltaK method predicted a weak modal signal at K=3, but the highest likelihood value occurred at K=1. Bar plots for K=2–4 showed a high degree of admixture among individuals across the five populations, consistent with high levels of gene flow across Australia (Figure 4, right panel). Pairwise F_st_ was then calculated for the five *P. australiana* populations using 976 SNPs. The global estimate was not significantly different from zero, indicating the populations are not differentiated (F_st_ = 0.0012, 99 % CI=0.0255–0.0403). Further, pairwise F_st_ values were low, ranging from -0.0033 to 0.0051, suggesting substantial gene flow among populations separated by distances between 341–2700 kilometres (Table 4).

**Table 4:** Pairwise comparisons of Weir and Cockerham’s (1984) *F_ST_* (below diagonal) and geographic distance in kilometres (above diagonal) among populations of *P. australiana* from five locations.

|  | Boomi | Calca | Esperance | Gilgandra | Goulburn |
| --- | --- | --- | --- | --- | --- |
| Boomi | — | 1555 | 2714 | 341 | 677 |
| Calca | -0.0033 | — | 1167 | 1365 | 1434 |
| Esperance | 0.0051 | 0.0028 | — | 2531 | 2572 |
| Gilgandra | 0.0001 | 0.0045 | 0.0000 | — | 364 |
| Goulburn | -0.0015 | -0.0007 | 0.0048 | 0.0003 | — |

### Insecticide susceptibility

Bioassays revealed highly contrasting responses to insecticide exposure in *P. xylostella* and *P. australiana* field strains (Figure 6). *Plutella australiana* showed extremely high susceptibility to all four insecticides evaluated: resistance ratios at the LC_50_ and LC_99_ estimates were less than 1.0, indicating that this strain was 1.5-fold to 7.4-fold more susceptible even than the laboratory *P. xylostella* (S) reference. In contrast, resistance ratios at the LC_50_ for the field *P. xylostella* strain ranged from 2.9 for Success Neo to 41.4 for Dominex (Table 5), indicating increased tolerance to all insecticides. The ratio of commercial field doses to LC_99_ estimates for each insecticide implied differences in field control between species. The field dose ratios for *P. xylostella* were between 0.1 for Dominex, indicating that a commercial dose of Dominex would fail to control this field strain, to 14.7 for Success Neo, and for *P. australiana* were between 17.7 for Dominex to 474.6 for Success Neo. Control mortality was similar for the field and reference strains, averaging 3.1% to 4.4% across all bioassays.

**Figure 6:**
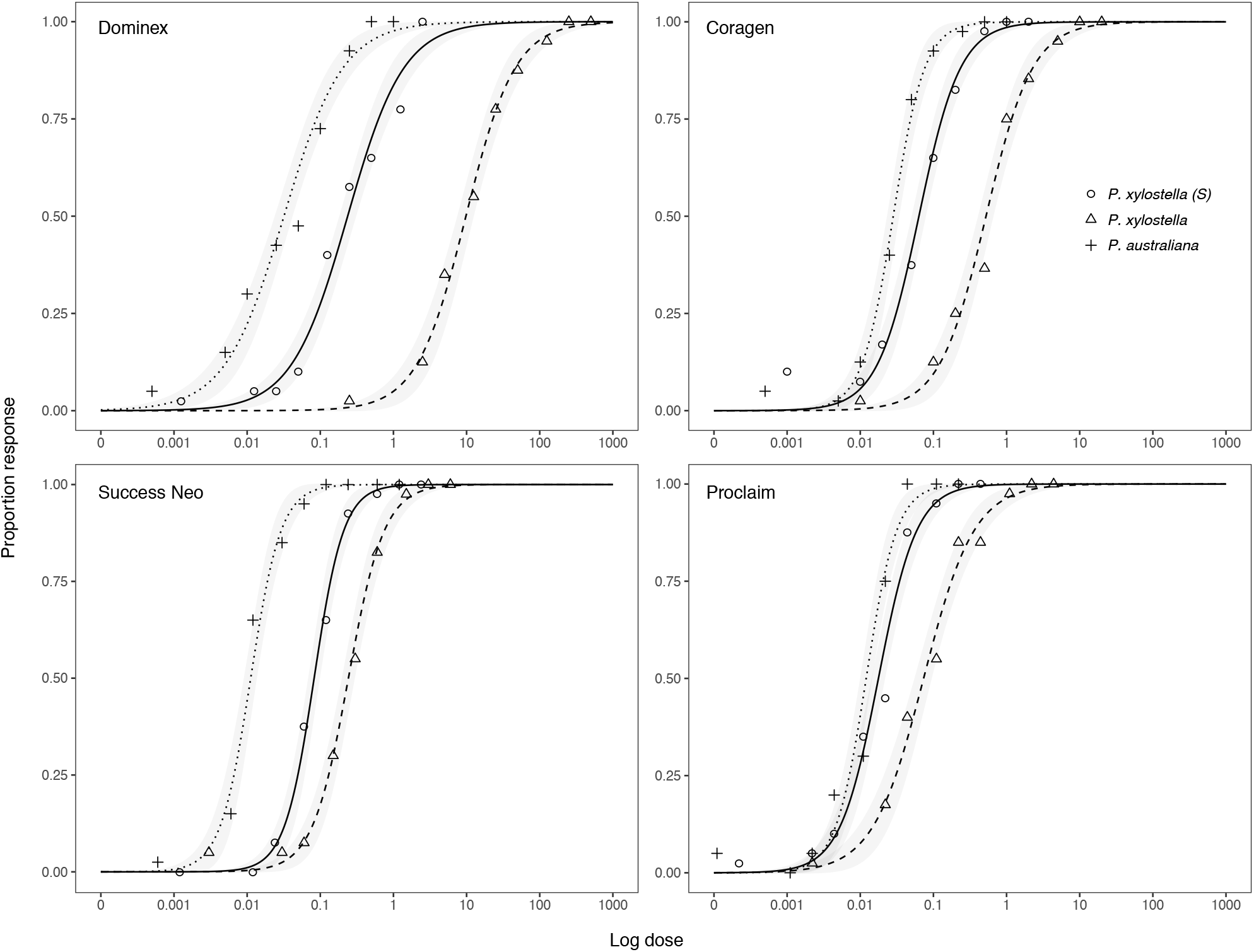
Dose response curves for *P. xylostella* and *P. australiana* field strains collected from Angle Vale and Urrbrae, South Australia, and a susceptible *P. xylostella* (S) reference strain, exposed to four commercial insecticides: Dominex, Coragen, Proclaim and Success Neo. Points are the mean observed response across four bioassay replicates and lines are the fitted log-logistic response curves with 95% confidence intervals shown in grey shading.

**Table 5:** Log-logistic regression statistics for dose-response bioassays on *P. australiana* (*P. aus*) and *P. xylostella* (*P.x*) field strains and the *P. xylostella* (*S*) reference strain exposed to four commercial insecticides. Statistics presented include the number of insects tested (*n*), *LC*_50_ and *LC*_99_ estimates with 95% confidence limits, resistance ratios (*RR*) at each *LC* level, and ratios of the commercial field doses to the *LC*_99_ estimates (Field dose ratio).

| Product | Strain | *n* | Slope (SE) | $LC_{50}$ (95% CL)<br>[mg L<sup>-1</sup> a.i.] | $RR_{LC50}$ | $LC_{99}$ (95% CL)<br>[mg L<sup>-1</sup> a.i.] | $RR_{LC99}$ | Field dose<br>ratio |
| --- | --- | --- | --- | --- | --- | --- | --- | --- |
| Coragen | *P. aus* | 320 | 2.016 ± 0.236 | 0.028 (0.023–0.034) | 0.45 | 0.276 (0.161–0.474) | 0.22 | 72.53 |
|  | *P. x* | 322 | 1.363 ± 0.149 | 0.524 (0.411–0.667) | 8.26 | 15.235 (7.374–31.479) | 11.88 | 1.31 |
|  | *P. x* (S) | 323 | 1.528 ± 0.165 | 0.063 (0.051–0.079) | 1.00 | 1.282 (0.666–2.47) | 1.00 | 15.60 |
| Dominex | *P. aus* | 320 | 1.078 ± 0.117 | 0.032 (0.024–0.042) | 0.13 | 2.267 (0.92–5.583) | 0.16 | 17.65 |
|  | *P. x* | 320 | 1.292 ± 0.146 | 9.792 (7.563–12.679) | 41.38 | 343.317 (158.25–744.816) | 24.85 | 0.12 |
|  | *P. x* (S) | 320 | 1.130 ± 0.118 | 0.237 (0.182–0.308) | 1.00 | 13.815 (5.685–33.574) | 1.00 | 2.90 |
| Proclaim | *P. aus* | 320 | 2.073 ± 0.235 | 0.012 (0.01–0.015) | 0.68 | 0.111 (0.066–0.186) | 0.39 | 119.40 |
|  | *P. x* | 320 | 1.254 ± 0.146 | 0.073 (0.056–0.096) | 4.15 | 2.868 (1.282–6.415) | 10.04 | 4.60 |
|  | *P. x* (S) | 320 | 1.652 ± 0.181 | 0.018 (0.014–0.022) | 1.00 | 0.286 (0.153–0.532) | 1.00 | 46.20 |
| Success Neo | *P. aus* | 320 | 2.087 ± 0.293 | 0.011 (0.009–0.014) | 0.14 | 0.101 (0.056–0.184) | 0.14 | 474.63 |
|  | *P. x* | 320 | 1.766 ± 0.196 | 0.242 (0.197–0.297) | 2.94 | 3.266 (1.805–5.912) | 4.65 | 14.69 |
|  | *P. x* (S) | 321 | 2.143 ± 0.255 | 0.082 (0.068–0.099) | 1.00 | 0.703 (0.417–1.184) | 1.00 | 68.28 |

**Table 6:**
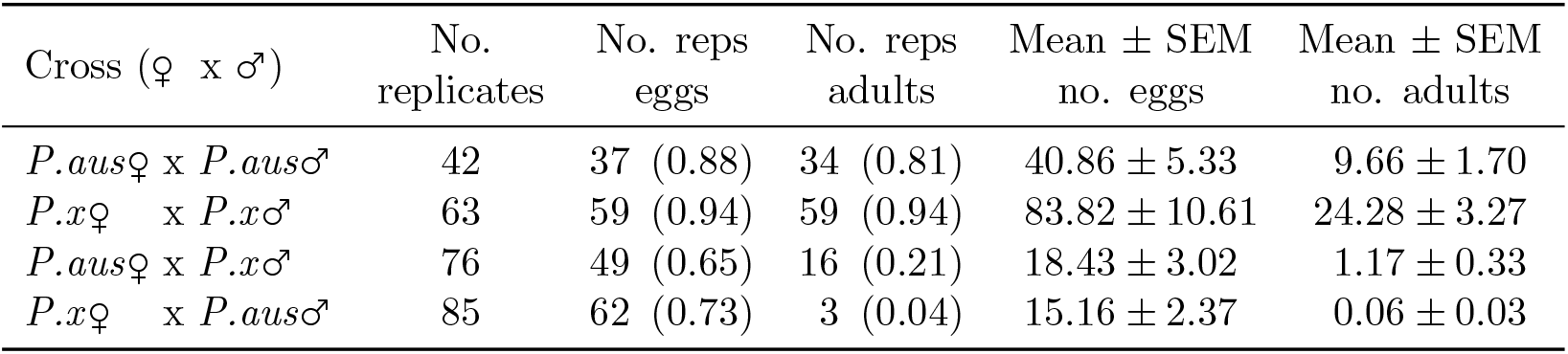
Fecundity of intra-species and reciprocal inter-species single pair crosses of *P. australiana* (*P.aus*) and *P. xylostella* (*P.x*). Presented are the number and proportion in parentheses of replicates that produced eggs and adult offspring, and the mean ± standard error of the mean number of eggs and adult offspring per replicate.

| Cross (♀ x ♂) | No.<br>replicates | No. reps<br>eggs | No. reps<br>adults | Mean ± SEM<br>no. eggs | Mean ± SEM<br>no. adults |
| --- | --- | --- | --- | --- | --- |
| *P. aus*♀ x *P. aus*♂ | 42 | 37 (0.88) | 34 (0.81) | 40.86 ± 5.33 | 9.66 ± 1.70 |
| *P.x*♀ x *P.x*♂ | 63 | 59 (0.94) | 59 (0.94) | 83.82 ± 10.61 | 24.28 ± 3.27 |
| *P. aus*♀ x *P.x*♂ | 76 | 49 (0.65) | 16 (0.21) | 18.43 ± 3.02 | 1.17 ± 0.33 |
| *P.x*♀ x *P. aus*♂ | 85 | 62 (0.73) | 3 (0.04) | 15.16 ± 2.37 | 0.06 ± 0.03 |

**Table 7:**
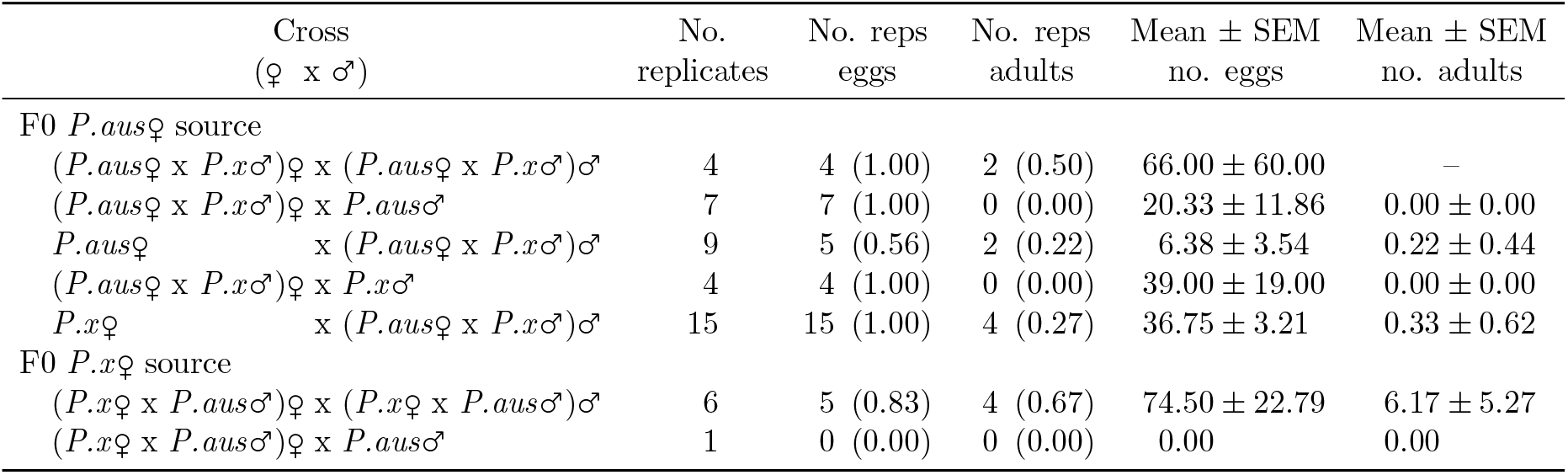
Fecundity of hybrid F1 crosses and back-crosses. Presented are the numbers and proportion in parentheses of replicates producing eggs and adult offspring, and the mean ± standard error of the mean numbers of eggs and adults offspring per replicate. A dash denotes an absence of count data.

## Discussion

Cryptic species arise when divergence does not lead to morphological change (Bickford et al., 2007). The recent discovery of a cryptic ally to the diamondback moth, *P. australiana*, was unexpected given the breadth of previous genetic studies on this economically important pest. Several factors may have contributed to this discovery, including the use of light traps for specimen collection, rather than limiting sampling to Brassica vegetable farms. Landry and Hebert (2013) also isolated DNA from legs, keeping most of each specimen intact and providing a morphological reference for examining unexpected genotypes. It is also possible that *P. australiana* was previously overlooked from nuclear DNA studies due to biases in amplification of divergent alleles. Here we sought to determine whether *P. australiana* is an agricultural pest, and to understand its ecological and genetic differences from *P. xylostella*.

Extensive larval sampling from wild and cultivated brassicaceous plants revealed that *P. australiana* widely co-occurs with *P. xylostella* throughout southern Australia, and utilizes some of the same host plants. The relative abundance of *P. australiana* was on average 9-fold lower than *P. xylostella*. We observed higher proportions of *P. australiana* in larval collections from the eastern state of New South Wales, similar to the light trap samples from Landry and Hebert (2013), possibly reflecting habitat suitability. Although we did not detect *P. australiana* in limited sampling from the island state of Tasmania, the presence of brassicas in the region and evidence from light traps that wind currents can transport *Plutella* moths across Bass Strait (Lionel Hill, Pers. Comm.) suggest it is likely to occur there.

Our study confirms that the host range of *P. australiana* includes canola crops and wild brassicaceous species. In laboratory rearing, *P. australiana* completed development on sand rocket, *D. tenuifolia*, and canola, Brassica napus, and was additionally collected from several other wild species without rearing to confirm host status. Our sampling focused on relatively few introduced brassicaceous species common in agricultural areas, yet the Australian Brassicales is represented by 11 plant families (Australasian Virtual Herbarium, https://avh.chah.org.au/). These include several families outside the Brassicaceae on which *P. xylostella* and allies have been documented feeding, including Capparaceae (Robinson & Sattler, 2001), Cleomaceae (Landry & Hebert, 2013) and Tropaeolaceae (Sarfraz et al., 2006). The Australian Brassicaceae has records for 61 genera and 205 species, including many introduced species but also a diversity of native genera, such as *Lepidium, Blennodium*, and *Arabidella*, that occur over vast areas of Australia. As *P. australiana* is apparently native, wider sampling of native Brassicales may identify other suitable hosts.

*Plutella australiana* larvae were not identified among samples from sixteen commercial brassica vegetable crops despite the high suitability of these crops for *P. xylostella* (Talekar & Shelton, 1993), however eggs were collected from Kale. It is possible that extreme insecticide susceptibility prevents juvenile *P. australiana* populations from establishing, as commercial vegetable crops are typically sprayed multiple times per crop cycle (Baker, 2011). Comparison of commercial field rates against LC_99_ estimates for the four evaluated insecticides suggest the likelihood of poor or marginal field control for some insecticides against *P. xylostella*, consistent with known levels of resistance (Baker, 2011; Endersby et al., 2008), but very high field control if the same rates were used against *P. australiana*. Alternatively, some vegetable cultivars may not be attractive for oviposition or suitable for larval survival in this species. Exposure to host plants stimulates reproductive behaviour in *P. xylostella* (Pivnick, Jarvis, Gillott, Slater, & Underhill, 1990). We noted that *P. australiana* cultures provided with cabbage seedlings failed to produce viable eggs over seven days, but after then replacing cabbage with Diplotaxis seedlings, egg-laying occurred within 24 hours. Olfactory cues for host recognition or oviposition (Justus & Mitchell, 1996; Renwick, Haribal, Gouinguene, & Stadler, 2006; J. Y. Sun, Sonderby, Halkier, Jander, & de Vos, 2009) may differ between *Plutella* species. Host preference and performance studies are required to test these hypotheses.

Insecticide bioassays have been routinely conducted on Australian *P. xylostella* to monitor levels of insecticide resistance in field populations (Baker, 2011; Endersby et al., 2008). This method appears unlikely to be affected by the presence of *P. australiana* under typical conditions, as a period of laboratory rearing is usually necessary to multiply individuals prior to screening. In our experience, laboratory rearing of the two *Plutella* species on cabbage plants selects against *P. australiana* individuals when competing with *P. xylostella* in cages, causing the complete loss of *P. australiana* within a few generations. The reasons for this are unknown but may include differences in host preference or development rate, or direct competition.

Crossing experiments revealed that hybridization can occur between *P. australiana* and *P. xylostella* under controlled conditions and is most likely to occur in crosses involving Wol-bachia-infected *P. australiana* females. Hybridization occurs in around 10% of animal species, particularly in captivity (Mallet, 2005), but asymmetric reproductive isolation is commonly observed in reciprocal crosses between taxa (Turelli & Moyle, 2007). In our experiments, a strong male bias in the offspring of interspecific crosses and failure to back-cross hybrid females both follow Haldane’s rule (Haldane, 1922), which predicts greater hybrid inviability or sterility in the heterogametic sex (female, in Lepidoptera). This pattern can arise from epistatic interactions between sex-linked and/or autosomal genes that result in genetic incompatibilities (C. Jiggins et al., 2001; Turelli & Orr, 2000). Although the back-crosses with F1 hybrid females were sterile, the back-crosses with hybrid males (to both species) were viable, which could enable the transfer of genes between hybrid and/or parental species. However, it is unclear whether hybridization occurs in the wild.

Although *P. australiana* and *P. xylostella* show deep divergence (8.6%) in mtDNA (Landry & Hebert, 2013), the sole use of mtDNA can be unreliable for inference of evolutionary history and should be corroborated using evidence from nuclear markers (Hurst & Jiggins, 2005). Our analysis revealed striking differences in nuclear diversity across the genome between co-existing populations of each *Plutella* species collected at the same locations and times, and from the same host plant species. *Plutella xylostella* populations from Australia and New Zealand have low levels of genetic diversity compared with populations from other continents, thought to reflect the recent introduction of this species from a small founding population (Endersby et al., 2006; Saw et al., 2006). Consistent with this view, we found a remarkable 1.5-fold reduction in heterozygosity across >300 000 sites in *P. xylostella* compared with sympatric *P. australiana* populations. However, both species showed limited mtDNA diversity with a single haplotype predominant. While outgroups from other continents were not available, comparative analysis of these closely-related Australian *Plutella* species suggested that patterns of mitochondrial and nuclear diversity are concordant in *P. xylostella* and consistent with a demographic bottleneck (Delgado & Cook, 2009; Saw et al., 2006), but discordant in *P. australiana*.

Mitochondrial variation can be strongly influenced by *Wolbachia* infection (Shoemaker et al., 2004). Extensive *Wolbachia* screening showed that each *Plutella* species was infected with a different strain at contrasting frequencies, and fit a ‘most-or-few’ pattern whereby species infection rates are often very low (<10%) or very high (>90%) (Hilgenboecker, Hammerstein, Schlattmann, Telschow, & Werren, 2008). Infection incidence in *P. xylostella* was lower in Australia (1%) than previously reported across global samples (5%) (Delgado & Cook, 2009). Our finding of a single supergroup A strain showing 100% sequence similarity to a strain reported in *P. xylostella* from Malaysia, plutWA1 (Delgado & Cook, 2009), provides some support of an Asian origin for Australian *P. xylostella* (Saw et al., 2006), though does not preclude this strain also occurring elsewhere.

Fixation of infection in *P. australiana* suggests that *Wolbachia* manipulates the reproductive biology of this species, though the host phenotype is unknown. We found no evidence of sex-ratio distortion, which has been associated with a *Wolbachia* strain, plutWB1, in *P. xylostella* (Delgado & Cook, 2009). High infection can be driven by cytoplasmic incompatibility (CI) (F. M. Jiggins, 2017). The high frequency of a single mtDNA haplotype among *P. australiana* individuals (87%) implies that the spread of *Wolbachia* infection has driven a selective sweep of co-inherited mtDNA through the population, causing a loss of mtDNA diversity (Shoemaker et al., 2004). High nuclear diversity (relative to sympatric *P. xylostella*) supports this hypothesis, because a demographic bottleneck should reduce diversity across the entire genome (Hurst & Jiggins, 2005). We found that *P. australiana* was infected with a supergroup B strain also reported to infect a mosquito, Culex pipiens, and a moth, Operophtera brumata (Derks et al., 2015; Havill et al., 2017). *Wolbachia* and host species often show incongruent phylogenies due to horizontal transfer of infections between taxa (Werren et al., 2008). Even closely related species often have different *Wolbachia* strains, demonstrating that most infections do not survive host speciation (Werren et al., 2008).

*Plutella australiana* and *P. xylostella* have co-existed in Australia for at least 125 years (>1300 generations), yet have strongly divergent mitochondrial and nuclear genomes, Wol-bachia infections and insecticide susceptibility phenotypes. Laboratory rearing and crossing experiments also suggested that interspecific differences in host plant use may exist. What explains such strong divergence between the two *Plutella* species, given sympatry and the capacity to hybridize? Endemism of *P. australiana* (Landry & Hebert, 2013) implies an ancient evolutionary history in Australia, and our data provide support for existing views that Australian *P. xylostella* were recently introduced from a small ancestral source population, possibly from Asia (Delgado & Cook, 2009; Juric et al., 2017; Saw et al., 2006). Therefore, the two *Plutella* species may have diverged in allopatry and recently come into secondary contact. Maintenance of divergence suggests strong continuing reproductive isolation, which can evolve as a side-effect of allopatric divergence (Telschow et al., 2014). All 100 individuals that were RAD sequenced showed concordance in nuclear multilocus genotypes and mtDNA genotypes identified through PCR-RFLP regardless of geographic location, as shown by structure analysis. Cryptic species in sympatry provides strong evidence of limited genetic exchange (Bickford et al., 2007). A small degree of genotypic admixture evident for a few individuals in the structure plots might be explained by ancestral polymorphism or some introgressive hybridization (Hedrick, 2013), or alternatively, could be an artefact if the dataset is not representative of the entire genetic background (Dupont et al., 2016). The level of hybridization that may be occurring between these species is unknown. Reproductive isolation may not be uniform across the genome (Harrison & Larson, 2014, 2016), and scans of larger genomic regions may be required to identify introgression and detect hybrids.

The factors leading to reproductive isolation between the two *Plutella* species in nature are unknown but could include a range of pre- or post-mating isolation mechanisms, such as as-sortive mating or hybrid fitness costs. Behavioural mating choices are often the main isolating factor in sympatric animals (Mallet, 2005). Does *Wolbachia* cause a reproductive barrier? The contrast in infection status creates the potential for cytoplasmic incompatibility between species (Jaenike, Dyer, Cornish, & Minhas, 2006). Interspecific crosses showed a pattern of asymmetric isolation consistent with the expected effects of unidirectional CI, where 21% crosses involving infected *P. australiana* females produced viable offspring, while the reciprocal CI-cross direction (uninfected *P. xylostella* males crossed with infected *P. australiana* males) was nearly sterile. However, this pattern was not continued in the F1 generation: infected hybrid males (derived from the *P. australiana* maternal line) produced offspring at comparable rates when back-crossed to either uninfected *P. xylostella* or infected *P. australiana* female parents. The role of *Wolbachia*-induced postzygotic isolation between the two *Plutella* species requires further study, however our results suggest it could be more important in the F0 generation. *Wolbachia* can contribute to post-zygotic genetic isolation after speciation by complementing hybrid incompatibilities (Gebiola, Kelly, Hammerstein, Giorgini, & Hunter, 2016; Jaenike et al., 2006). Symbiont infections could also influence mating behaviour and contribute to pre-mating isolation (Shropshire & Bordenstein, 2016).

## Conclusions

The discovery of cryptic pest species introduces complexities for their management and also exciting opportunities for understanding ecological traits. We found strong genomic and phenotypic divergence in two cryptic *Plutella* lineages co-existing in nature, supporting their status as distinct species (Landry & Hebert, 2013), despite their capacity to hybridize. Reproductive isolation is likely to have evolved during allopatric speciation, and genome-wide sequence data suggest it has been maintained following secondary contact. Variation in *Wolbachia* infections might be one factor reinforcing reproductive barriers.

*Plutella australiana* co-occurs with *P. xylostella* throughout agricultural regions of southern Australia, but made up only 10% of *Plutella* juveniles collected from cultivated and wild brassicaceous plants. A lack of population structure across neutral SNP markers suggests that *P. australiana* populations are linked by high levels of gene flow, which is supported by light trap collections (Landry & Hebert, 2013) and seasonal colonization of canola crops. Future molecular analysis of Australian *Plutella* should include a species identification step using PCR-RFLP. For ecological studies, it may be possible to perform molecular species identification to confidently distinguish a representative sub-sample of individuals or pooled samples. Our study has shown that while *P. australiana* can attack canola crops, there is no evidence of pest status in commercial brassica vegetables, and bioassays suggested that field populations should be easily controlled with insecticides. Though *P. australiana* is a potential pest of some Australian Brassica crops, it is of secondary importance to the diamondback moth, *P. xylostella*.

## Acknowledgements

We thank the many colleagues who collected *Plutella* samples: Adam Hancock, Adam Pearce, Adam Quade, Alan Lord, Andrew McMahen, Andy Bates, Andy Ryland, Brenton Spriggs, Chris Davey, Chris Teague, Craig James, Dustin Berryman, Grant Hudson, Guy Westmore, James McKee, Jessica Smith, Jo Holloway, Joanne Holloway, Josh Andrews, Josh Hollitt, Karina Bennett, Laura Archer, Levon Cookson, Lisa Ohlson, Louise Flohr, Melina Miles, Michael Collins, Monica Field, Nigel Myers, Peter Cole, Peter Ellison, Peter Gregg, Peter Mangano, Richard Saunders, Sarina Macfadyen, Stewart Learmonth. We also thank Lynn Dieckow and Birte Albrecht for technical assistance with genotyping. KDP was supported by the University of Adelaide (UA00146) and the Grains Research and Development Corporation (GRDC) (DAS00094), GJB, KJP and JKK were supported by GRDC (DAS00155), and SWB was supported by the Australian Research Council (DP120100047, FT140101303).

## Data accessibility

DNA sequences: GenBank accession numbers MF804301-MF804314 (wsp) and MF151826-MF151906 (COI). RADseq FASTQ files will be submitted to the NCBI Sequence Read Archive.

## Author contributions

Wrote the manuscript: KDP

SWB Conceived and designed experiments: All

Data analysis: KDP, SWB, CMW

Sample collection: KDP

COI genotyping, RADseq: KDP, SWB

*Wolbachia* genotyping: SWB, CMW

*Plutella* cultures, bioassays and crossing experiments: KJP, JKK, GJB

